# Proteinase 3 is involved in presepsin production through neutrophil extracellular trap phagocytosis by macrophages

**DOI:** 10.1101/2025.05.02.651854

**Authors:** Akishige Ikegame, Akihiro Kondo, Tatsuya Morinishi, Yusuke Yamaguchi, Yui Okutani, Chinatsu Maegawa, Satoshi Tada

## Abstract

Presepsin (P-SEP), a soluble subtype of CD14, has been widely reported as a useful biomarker for sepsis in clinical studies. However, the cytological mechanism underlying its production remains poorly understood. Previous research has demonstrated that P-SEP is generated when M1 macrophages (M1 MΦs) phagocytose neutrophil extracellular traps (NETs) and degrade internalized CD14. In this study, we demonstrated that M1 MΦs phagocytose NETs and that proteinase 3 (PR3) within M1 MΦs degrades NET-derived CD14, resulting in the production of P-SEP. Furthermore, we observed a strong correlation between the amount of NETs and P-SEP levels, indicating that NET phagocytosis is a key process in P-SEP generation. Our findings also suggest that PR3 released extracellularly along with NETs may degrade CD14 in plasma, contributing to elevated P-SEP levels. This mechanism may contribute to increased P-SEP concentrations observed in autoimmune diseases associated with NET accumulation, proposing P-SEP as a potential biomarker of disease activity beyond sepsis.

## Introduction

Presepsin (P-SEP) is currently used in clinical practice as a biomarker for sepsis. However, elevated P-SEP levels have also been reported in patients with autoimmune diseases, such as systemic lupus erythematosus and hemophagocytic syndrome, even in the absence of infection; however, the underlying mechanisms remain unclear [1–3]. If the cytological mechanism of P-SEP production can be elucidated, it may pave the way for its development as a biomarker not only for sepsis but also for autoimmune diseases.

CD14, the precursor of P-SEP, is a 55 kDa glycosylphosphatidylinositol (GPI)-anchored glycoprotein that acts as a key pattern recognition receptor in the innate immune system. When lipopolysaccharide (LPS) released from bloodborne bacteria binds to CD14 and myeloid differentiation protein 2 (MD2) on the cell surface, it forms the LPS–CD14–MD2 complex. This complex activates the nuclear factor-kappa B (NF-κB) signaling pathway via binding to toll-like receptor 4 (TLR4), thereby initiating an inflammatory response [4–6]. CD14 is expressed on the surface of monocytes (Mos), macrophages (MΦs), and neutrophils, and also exists in soluble form in the serum. Increased CD14 expression has been identified as a biological risk factor for the development of sepsis and subsequent organ dysfunction in both animal models and clinical studies [7,8].

P-SEP is a 13 kDa glycoprotein released extracellularly as a cleavage product of the N-terminal portion of CD14. Under conventional understanding, P-SEP is produced when CD14 expressed on the surface of Mos or MΦs is internalized together with phagocytosed bacteria and subsequently degraded by the lysosomal protease cathepsin D [9,10]. Arai et al. reported that monocyte phagocytosis contributes more significantly to P-SEP production than neutrophil phagocytosis, with cathepsin D and elastase mediating CD14 degradation in monocytes [11].

In our prior investigations into the mechanisms of P-SEP production, we focused on neutrophil extracellular traps (NETs), which are known to increase in the peripheral blood during sepsis and autoimmune diseases [12–14]. NETs are mesh-like structures composed of decondensed chromatin with pores approximately 200 nm in diameter. They are extruded by neutrophils as part of the host defense mechanism and are decorated with nuclear proteins (e.g., histones), granule proteins (e.g., neutrophil elastase and myeloperoxidase), and cytoplasmic proteins (e.g., S100A8, A9, A12, actin, and α-actinin) [15–17]. NET formation is triggered by pathogen-and damage-associated molecular patterns released from bacteria. Reactive oxygen species activate peptidyl arginine deiminase 4 (PAD4), which citrullinates histones and promotes chromatin decondensation, ultimately leading to NET release into the extracellular space [18,19].

We previously reported that phagocytosis of NETs by Mos or MΦs induces higher P-SEP production than does phagocytosis of bacteria alone [20]. Additionally, we demonstrated that Mos remain in the circulation only for a few hours, whereas M1 MΦs that migrate into tissues are the primary producers of P-SEP. Animal studies using murine sepsis models revealed that M1 MΦs residing in the lung, liver, and kidneys are major contributors of P-SEP production [21]. In the course of these studies, we found that inhibition of cathepsin D and elastase reduced—but did not completely eliminate—P-SEP production, suggesting the involvement of other proteolytic enzymes in CD14 degradation.

Proteinase 3 (PR3) is a serine protease found in neutrophils, Mos, and in the azurophilic granules of macrophages. It is primarily expressed intracellularly in neutrophils under non-inflammatory conditions, but translocates to the plasma membrane surface during inflammatory responses, including sepsis [22,23]. Extracellular PR3 has also been implicated in vascular dysfunction and increased vascular permeability [24]. Furthermore, PR3-expressing neutrophils, monocytes, and macrophages contribute to persistent inflammation by releasing chemokines, such as monocyte chemoattractant protein-1 (MCP-1), regulated on activation, normal T-cell expressed and secreted (RANTES), macrophage inflammatory protein (MIP)-1α, and MIP-1β, when phagocytosed [25,26]. PR3 also plays a role in MΦ polarization, promoting M1-type differentiation while suppressing M2 MΦs [27,28].

Most existing studies on PR3 have focused on its extracellular functions and its role in maintaining inflammation; however, the cytological function of PR3 within M1 MΦs has not been explored. In this study, we aimed to determine whether M1 MΦs phagocytose NETs and whether PR3 within these cells degrades NET-derived CD14, leading to P-SEP production.

## Materials and Methods

### Reagents and antibodies

Details of the reagents used in this study and their sources are as follows: Polymorphprep^™^ (Cat. No.1114683; Axis-Shield, Dundee, Scotland), RPMI-1640 Medium (Cat. No. R8758; Sigma-Aldrich, MO, USA), phorbol 12-myristate 13-acetate (PMA; Cat. No. P8139; Sigma-Aldrich, MO, USA), *Escherichia coli* DH5α competent cells (*E. coli* DH5α; Cat. No. 9057; Takara Bio, Shiga, Japan), EasySep^™^ Human Monocyte Isolation Kit (Cat. No. 19359; Stemcell Technologies, Vancouver, BC, Canada), CellXVivo^TM^ Human M1 Macrophage Differentiation Kit (Cat. No. CDK012; R&D systems, Minneapolis, MN, USA), IntraPrep Permeabilizaton Reagent (Cat. No. A07803; Beckman Coulter, Brea, CA, USA), cytochalasin D (Cat. No. 11330; Cayman Chemical, Ann Arbor, MI, USA), wortmannin (Cat. No. AG-CN2-0023-M001; Adipogen Life Sciences, San Diego, CA, USA), phenylmethylsulfonyl fluoride (PMSF; Cat. No. 195381; MP Biomedicals, Irvine, CA, USA), elafin (Cat. No. 4243-v; Peptide Institute, Osaka, Japan), May–Grünwald’s stain solution (Cat. No. 15053; Mutokagaku, Tokyo, Japan), Giemsa stain solution (Cat. No. 15002; Mutokagaku, Tokyo, Japan), 1/15 M phosphate buffer solution (pH 6.4) (Cat. No. 15612; Mutokagaku, Tokyo, Japan), 4% paraformaldehyde phosphate buffer solution (Cat. No. 09154-85; Nacalai Tesque, Kyoto, Japan), Triton^™^ X-100 (Cat. No. 194854; MP Biomedicais, Irvine, CA, USA), sample buffer solution with 2-mercaptoethanol (Cat. No. 30566-22; Nacalai Tesque, Kyoto, Japan), skim milk powder (Cat. No. 190-12865; Fujifilm Wako Pure Chemical, Tokyo, Japan), Chemi-Luna One Super (Cat. No. 02230; Nacalai Tesque, Kyoto, Japan), Cell Death Detection ELISAPLUS (Cat. No. 11774425001; Roche Diagnostics, Tokyo, Japan), SYTOX^™^ Green (Cat. No. S7020; Invitrogen, Waltham, MA, USA), Hoechest^®^ 33342 (Cat. No. 346-07951; Fujifilm, Tokyo, Japan), Recombinant human CD14 protein (rCD14; Cat. No. 383-CD; R&D systems, Minneapolis, MN, USA), and recombinant human PR3 protein (Cat. No. 13699-H08H1; Sino Biological, Kanagawa, Japan).

Details of the antibodies used in this study and their sources are as follows: FITC-conjugated anti-human PR3 mouse monoclonal antibody (Cat. No. ab65255; Abcam, Cambridge, UK), PE-conjugated anti-human cathepsin D mouse monoclonal antibody (Cat. No. sc-13148 PE; Santa Cruz Biotechnology, Dallas, TX, USA), FITC-conjugated anti-human neutrophil elastase mouse monoclonal antibody (Cat. No. sc-55549 FITC; Santa Cruz Biotechnology, Dallas, TX, USA), PE-conjugated anti-human flavocytochrome b558 mouse monoclonal antibody (Cat. No. D162-5; Medical & Biological Laboratories, Tokyo, Japan), PE/Cyanine7-conjugated anti-human CD14 mouse monoclonal antibody (Cat. No. 367111; BioLegend, San Diego, CA, USA), FITC-conjugated anti-human CD68 mouse monoclonal antibody (Cat. No. F7135; DAKO, Glostrup, Denmark), FITC-conjugated anti-human CD282 (TLR2) mouse monoclonal antibody (Cat. No. 309705; BioLegend, San Diego, CA, USA), PE-conjugated anti-human CD284 (TLR4) mouse monoclonal antibody (Cat. No. 312805; BioLegend, San Diego, CA, USA), anti-citrullinated histone H3 (Cit-H3) (citrulline R2 + R8 + R17) rabbit polyclonal antibody (Cat. No. ab5103; Abcam, Cambridge, UK), anti-human LL-37 mouse monoclonal antibody (Cat. No. sc-166770; Santa Cruz Biotechnology, Dallas, TX, USA), anti-β-actin rabbit monoclonal antibody (Cat. No. 4970; Cell Signaling Technology, Danvers, MA, USA), anti-human CD14 mouse monoclonal antibody (Cat. No. 14-0149-82; Invitrogen, Waltham, MA, USA), anti-PR3 mouse monoclonal antibody (Cat. No. sc-74534; Santa Cruz Biotechnology, Dallas, TX, USA), anti-human presepsin mouse monoclonal antibody (F1106-13–3; Mochida Pharmaceutical, Tokyo, Japan), anti-human presepsin rabbit monoclonal antibody (S68; Mochida Pharmaceutical, Tokyo, Japan), tetramethylrhodamine (TRITC)-conjugated goat anti-mouse IgG (H+L) secondary antibody (Cat. No. SA00007-1; Cosmo Bio, Tokyo, Japan), TRITC-conjugated goat anti-rabbit IgG (H+L) secondary antibody (Cat. No. SA00007-2, Cosmo Bio, Tokyo, Japan), Alexa Flour 405-conjugated goat anti-rabbit IgG (H+L) secondary antibody (Cat. No. ab175652; Invitrogen, Waltham, MA, USA), horseradish peroxidase (HRP)-conjugated goat anti-mouse IgG (H+L) secondary antibody (Cat. No. 7076; Cell Signaling Technology, Danvers, MA, USA), HRP-conjugated gort anti-rabbit IgG (H+L) secondary antibody (Cat. No. 7074; Cell Signaling Technology, Danvers, MA, USA).

### Purification of human neutrophils

Peripheral blood samples were collected from five healthy participants using ethylene diamine tertaacetic acid (EDTA)-2K-containing collection tubes. The anticoagulated blood was layered onto Polymorphprep™ at 20 ± 2 °C. Following density gradient centrifugation at 500 × *g* for 30 minutes at room temperature (RT, 20 ± 5 °C), the neutrophil-rich lower layer was harvested. Serum-free RPMI 1640 medium was added, and the cells were washed at 400 × *g* for 10 minutes to obtain purified neutrophils. Cell purity was confirmed to be ≥98.0% using a multiparameter automated hematology analyzer (XS-800i, Sysmex, Hyogo, Japan).

### Mos Isolation

Peripheral blood samples from five healthy participants were processed using EDTA-2K–containing tubes. Blood was layered onto Polymorphprep™, and the upper layer containing peripheral blood mononuclear cells (PBMCs) was collected after density gradient centrifugation. The PBMCs were washed twice with serum-free RPMI 1640 at 100 × *g* for 10 minutes to remove platelets. Mos were then isolated by negative selection using the EasySep™ Human Monocyte Isolation Kit. Cell purity was confirmed to be ≥94.0% using the XS-800i.

### Differentiation of Mos into M1 MΦs

Mos were differentiated into M1 MΦs using the CellXVivo™ Human M1 Macrophage Differentiation Kit. Purified Mos (2.0 × 10⁶ cells/well) were suspended in a medium containing 10 ng/mL granulocyte-macrophage colony-stimulating factor and cultured at 37 °C in a 5% CO₂ atmosphere for 6 days. After differentiation, the M1 MΦs were stained with May–Grünwald–Giemsa (MG) stain, and their morphology was observed at 1,000× magnification using a BX50 biological microscope (Olympus, Tokyo, Japan). In addition, the surface expression of CD14, CD45, CD68, CD80, CD86, and CD163 was evaluated by staining with fluorescently conjugated antibodies, followed by analysis of CD14⁺CD45⁺ M1 MΦs using a flow cytometer (CytoFLEX, Beckman Coulter, Brea, CA, USA).

### Comparison of the proteolytic enzyme levels in neutrophils, Mo, and M1 MΦ

Neutrophils and Mos isolated from the peripheral blood of healthy participants, as well as M1 MΦs differentiated from Mos, were used. After fixation and permeabilization of each cell type using IntraPrep, FITC-conjugated anti-PR3 antibody, PE-conjugated anti-cathepsin D antibody, and FITC-conjugated anti-elastase antibody were added and incubated at RT for 20 minutes in the dark. After washing with phosphate-buffered saline (PBS), the proteolytic enzyme levels were analyzed using CytoFLEX. Lymphocytes isolated from the peripheral blood of healthy participants were used as negative controls, and the proteolytic enzyme expression in each phagocyte was evaluated based on mean fluorescence intensity (MFI) using CytoFLEX.

### NET induction and analysis using flow cytometry

Purified neutrophils (2.0 × 10⁶ cells/well) were suspended in serum-free RPMI 1640 medium and seeded into 24-well plates (Cat. No. SIAL0526, Sigma-Aldrich, Burlington, MA, USA). NETs were induced by stimulation with either an *E. coli* DH5α suspension adjusted to an optical density (OD) of 1.0 at 660 nm, incubated at 37 °C in a 5% CO₂ atmosphere for 4 hours, or with 50 nM PMA under the same conditions for 2 hours. As described previously, NETs induced by *E. coli* DH5α and PMA were referred to as *E. coli* DH5α-NETs and PMA-NETs, respectively [21].

Following NET induction, cells were gently resuspended by pipetting to obtain a neutrophil suspension containing NETs. The suspension was stained with an anti-Cit-H3 antibody at RT for 20 minutes, followed by incubation with a TRITC-conjugated anti-rabbit IgG antibody at RT for 20 minutes. After staining, Cit-H3 positivity was assessed by flow cytometry using a CytoFLEX flow cytometer. Forward scatter (FS) and side scatter (SS) parameters were also analyzed. In accordance with previous studies, regions with >60% Cit-H3 positivity were defined as NET areas (Fig. 2a). In addition, we compared the fluorescence intensities of flavocytochrome b558, SYTOX™ Green, CD14, TLR2, TLR4, and LL-37 in the NET area after *E. coli* DH5α stimulation with those in the neutrophil area, defined as regions of untreated neutrophils (Fig. 2b).

### Evaluation of NET ratio by enzyme-linked immunosorbent assay

Untreated neutrophils and cell suspensions collected after stimulation with *E. coli* DH5α or PMA were harvested by gentle pipetting and centrifuged at 200 × *g* for 10 minutes. The resulting supernatants were used as samples for an enzyme-linked immunosorbent assay (ELISA), performed using the Cell Death Detection ELISAPLUS kit, which employs streptavidin-coated microplates. After adding the samples to the wells, DNA–histone complexes—characteristic components of NETs—were detected using a biotinylated anti-histone antibody and a peroxidase-conjugated anti-DNA antibody. Absorbance was measured at 405 nm using a microplate reader (Bio-Rad, Berkeley, CA, USA) with 2,2’-azino-bis(3-ethylbenzothiazoline-6-sulfonic acid) (ABTS) as the chromogenic substrate. The concentration of DNA–histone complexes in each sample was interpreted as the NET ratio (Fig. 2c).

### Detection of Cit-H3 and CD14 using western blotting

Untreated neutrophils and cell suspensions obtained after *E. coli* DH5α or PMA stimulation were collected and centrifuged at 15,000 × *g* for 5 minutes at 4 °C. The resulting cell pellets were resuspended in 100 μL of sample buffer containing 2-mercaptoethanol, boiled at 97 °C for 3 minutes, and stored at –80 °C until further analysis. Proteins were separated on a 15% polyacrylamide gel and transferred to a polyvinylidene fluoride (PVDF) membrane at 120 mA at RT for 60 minutes. The membrane was blocked with 5% skim milk at RT for 60 minutes, followed by incubation with anti-CD14 (1:1,000), anti-Cit-H3 (1:1,000), and anti-β-actin (1:15,000) antibodies at 4 °C for 12 hours.

After washing with tris-buffered saline containing 0.1% Tween 20 (TBST) for 10 minutes, three times, the membrane was incubated with HRP-conjugated anti-rabbit IgG antibody (1:2,500) at RT for 60 minutes. Following three additional washes with TBST, protein bands were detected using Chemi-Lumi One Super. Band intensities were quantified using ImageJ software (version 1.8.0; National Institutes of Health, Bethesda, MD, USA). Protein expression levels of Cit-H3 and CD14 were normalized to β-actin, which was used as an internal control (Fig. 2d, Supplementary Fig. 1).

### Observation of M1 MΦs that phagocytosed NET

M1 MΦs (5.0 × 10⁵ cells/well) were seeded onto coverslips placed in 24-well plates and co-cultured with NETs at 37 °C in a 5% CO₂ atmosphere for 180 minutes. After co-culture, the cells on the coverslips were stained with MG stain, and cell morphology was observed at 1,000× magnification using a BX50 biological microscope. The P-SEP level in the culture supernatant was measured using an immunoluminescence analyzer (PATHFAST, PHC Holdings, Tokyo, Japan). The correlation coefficient between P-SEP levels and NET ratio, as measured by ELISA, was then determined.

### P-SEP production in M1 MΦ phagocytosis of NET using immunofluorescence staining

M1 MΦs (5.0 × 10⁵ cells/well) were seeded onto coverslips placed in 24-well plates and co-cultured with NETs at 37 °C in a 5% CO₂ atmosphere for 60 minutes. After co-culture, the cells on the coverslips were fixed with 4% paraformaldehyde in PBS, washed with PBS, and blocked with 0.1% Triton™ X-100/PBS containing 5% rat serum for 60 minutes. The cells were then incubated with either anti-Cit-H3 antibody (1:1,000) or anti-P-SEP (F1106-13-3) antibody (1:2,000) for 60 minutes. After washing, the cells were further incubated with Alexa Fluor 405-conjugated anti-rabbit IgG antibody (1:1,000), TRITC-conjugated anti-mouse IgG antibody (1:5,000), or FITC-conjugated anti-CD68 antibody (1:1,000) for 90 minutes. All subsequent procedures after cell culture were performed at RT. After the final wash, the coverslips were mounted onto glass slides, sealed with 80% glycerol, and imaged using a confocal microscope (FLUOVIEW FV10i, Olympus, Tokyo, Japan).

### Effect of PR3 inhibition of M1 MΦs on P-SEP production

To evaluate the effect of PR3 inhibition on P-SEP production, M1 MΦs were preincubated with PR3 inhibitors and subsequently co-cultured with NETs. As a control, M1 MΦs co-cultured with neutrophils in the absence of inhibitors were used. For PR3 inhibition in M1 MΦs, the serine protease inhibitors PMSF and elafin were employed.

M1 MΦs (5.0 × 10⁵ cells/well) were treated with either 10 μM PMSF or 0.5 μM elafin at 37 °C in a 5% CO₂ atmosphere for 30 minutes prior to co-culture. They were then co-cultured with either untreated neutrophils or PMA-NETs under the same conditions for 180 minutes. After co-culture, the P-SEP concentration in the supernatants was measured using the PATHFAST immunoassay system.

In parallel, M1 MΦs co-cultured with PMA-NETs in the presence of PMSF or elafin were harvested 15 minutes after the start of co-culture, and P-SEP protein levels were assessed by western blotting (Supplementary Fig. 2).

### Detection of P-SEP produced by M1 MΦ after NET phagocytosis using western blotting

After co-culturing M1 MΦs with PMA-NETs at 37 °C in a 5% CO₂ atmosphere for 15 minutes, the entire cell suspension was collected. The cells were pelleted by centrifugation at 15,000 × *g* for 5 minutes at 4 °C. The resulting pellet was resuspended in 100 μL of sample buffer containing 2-mercaptoethanol, boiled at 97 °C for 3 minutes, and stored at –80 °C until further analysis. Proteins were separated on a 15% polyacrylamide gel and transferred to a PVDF membrane at 120 mA for 60 minutes at RT. The membrane was blocked with 5% skim milk at RT for 60 minutes, followed by incubation with anti-P-SEP (S68) antibody (1:500) and anti-β-actin antibody (1:15,000) at 4 °C for 12 hours. After three washes with TBST, the membrane was incubated with HRP-conjugated anti-rabbit IgG secondary antibody (1:2,500) at RT for 60 minutes. Following another three washes with TBST, P-SEP and β-actin bands were visualized using Chemi-Lumi One Super, and the amount of P-SEP protein was quantified. β-actin was used as an internal control, and the P-SEP band intensity was normalized to that of β-actin.

### Evaluation of P-SEP production by inhibition of NET phagocytosis in M1 MΦ

To evaluate the effect of NET phagocytosis inhibition on P-SEP production, M1 MΦs were treated with cytochalasin D and wortmannin. M1 MΦs (5.0 × 10⁵ cells/well) were incubated with 50 μM cytochalasin D or 10 μg/mL wortmannin at 37 °C in a 5% CO₂ atmosphere for 30 minutes prior to co-culture. Cytochalasin D and wortmannin were used to inhibit NET phagocytosis by M1 MΦs. Prior to co-culture with NET, M1 MΦs (5.0 × 10⁵ cells/well) were treated with 50 μM cytochalasin D or 10 μg/mL wortmannin at 37 °C for 30 minutes. The optimal concentration of wortmannin was determined to be 10 μg/mL based on dose–response experiments using concentrations of 0.1, 1.0, and 10 μg/mL (Supplementary Fig. 3). P-SEP levels in the culture supernatants were measured using the PATHFAST.

In parallel, M1 MΦs treated with each inhibitor and NETs were collected 15 minutes after co-culture, and P-SEP protein levels were evaluated by western blotting (Supplementary Fig. 4).

### Evaluation of P-SEP production by PR3 using recombinant human CD14

rCD14 was dissolved in PBS to a final concentration of 50 ng/mL and incubated with 10 μg/mL PR3 at RT for 60 minutes. To inhibit PR3-mediated degradation of CD14, 10 μM PMSF or 0.5 μM elafin was added to the rCD14 solution immediately after PR3 addition. P-SEP levels in the supernatants of each sample were measured using the PATHFAST.

### Detection of recombinant human CD14 protein and produced P-SEP using western blotting

PR3 was added to the rCD14 solution, and 100 μL samples were collected after 60 minutes. Similarly, 100 μL samples were collected following PR3 inhibition with 10 μM PMSF or 0.5 μM elafin. Each 100 μL sample was mixed with 100 μL of 2× sample buffer containing 2-mercaptoethanol, boiled at 97 °C for 3 minutes, and stored at –80 °C until further analysis. Proteins were separated by SDS-PAGE on a 15% polyacrylamide gel and transferred to a PVDF membrane at 120 mA for 60 minutes at RT. The membrane was blocked with 5% skim milk at RT for 60 minutes, followed by incubation with anti-CD14 antibody (1:1,000) and anti-P-SEP (S68) antibody (1:500) at 4 °C for 12 hours. After washing three times with TBST, the membrane was incubated with HRP-conjugated anti-rabbit IgG secondary antibody. Following three additional washes with TBST, CD14 and P-SEP bands were visualized using Chemi-Lumi One Super. Band intensities were quantified using ImageJ to determine the amount of P-SEP protein (Supplementary Fig. 5).

### Statistical analyses

All data are expressed as mean ± standard deviation (SD). Statistical analyses were performed using JMP Pro 17 (SAS Institute, Cary, NC, USA). One-way analysis of variance (ANOVA) was conducted, and a *p*-value < 0.05 was considered statistically significant. When significant differences were observed among the groups, Dunnett’s test was used for comparisons with the control group, and the Tukey–Kramer honestly significant difference (HSD) test was applied for all pairwise comparisons. In the figures, * indicates *p* < 0.05, and ** indicates *p* < 0.01.

## Ethics statement

This study was conducted per the principles of the Declaration of Helsinki and was approved by the Ethical Review Committee of Kagawa Prefectural University of Health Sciences (Approval No. 443). Written informed consent was obtained from all the participants for blood collection and subsequent analyses.

## Results

### M1 MΦ cells contained high amounts of PR3

Flow cytometric analysis was performed to compare the intracellular proteolytic enzyme levels among cell types using MFI values. The MFI of PR3 in lymphocytes (negative control) was 4,726 ± 234.7, whereas markedly higher values were observed in M1 MΦ (43,517 ± 812.1), Mos (31,520 ± 391.3), and neutrophils (30,285 ± 145.3), indicating that PR3 was highly expressed in all the phagocytic cells. Similarly, the MFIs of cathepsin D and elastase in lymphocytes were 731 ± 40.1 and 897 ± 47.5, respectively. In M1 MΦ, the MFIs were 13,683 ± 106.3 for cathepsin D and 12,415 ± 333.5 for elastase; in Mos, 1,697 ± 63.0 and 2,223 ± 5.7; and in neutrophils, 2,092 ± 12.5 and 2,894 ± 10.7, respectively. These results demonstrate that M1 MΦ contain the highest levels of the proteolytic enzymes PR3, cathepsin D, and elastase among the analyzed cell types (Fig. 1).

**Fig 1.**
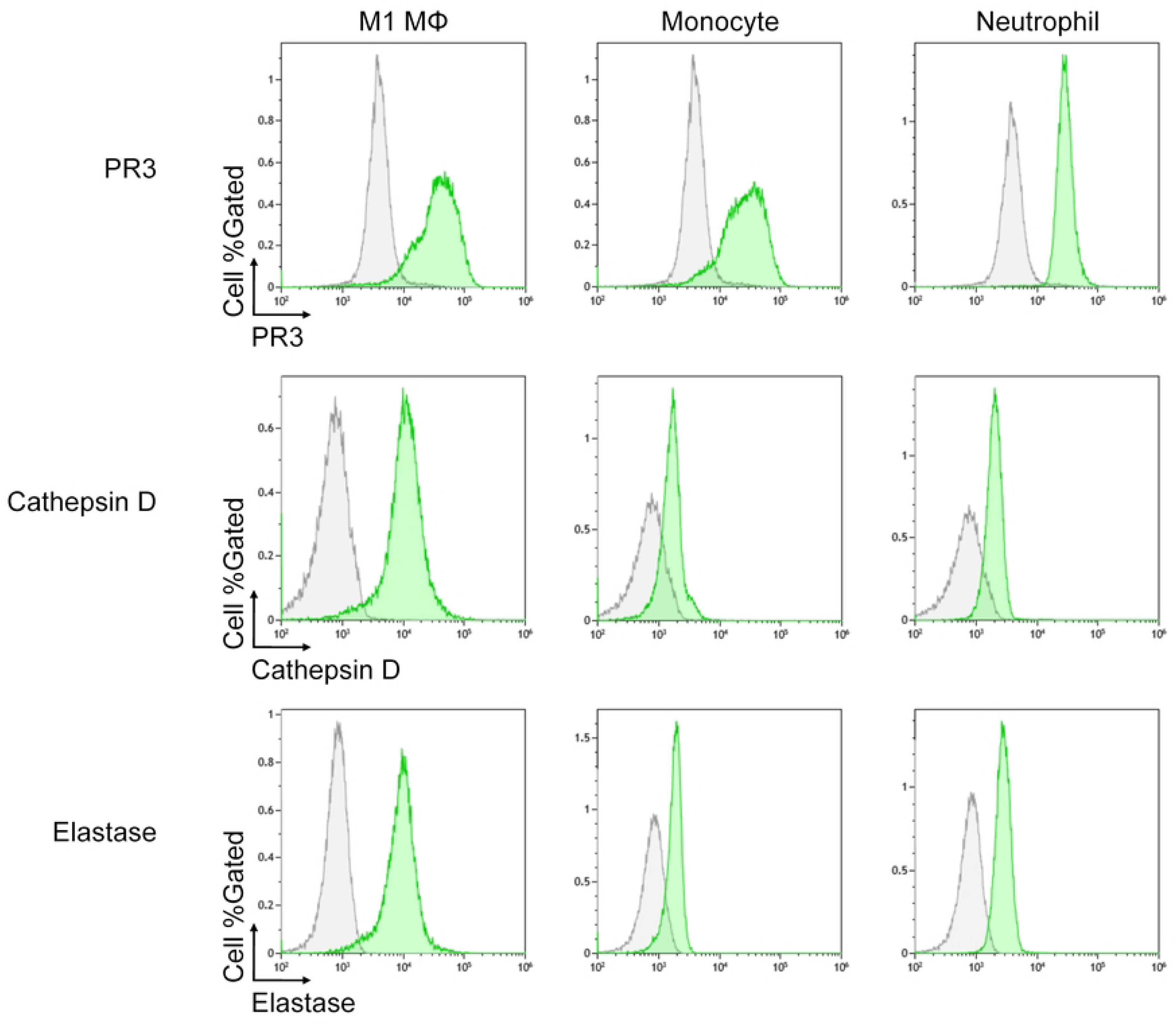
Comparison of the proteolytic enzyme levels in phagocytes. Three types of blood-derived cells (neutrophils and Mos isolated from the peripheral blood of healthy participants, and M1 MΦ differentiated from Mos) were analyzed for intracellular levels of PR3, cathepsin D, and elastase by flow cytometry. Proteolytic enzyme levels were compared using MFI. Lymphocytes were included as negative controls and are shown in gray, while phagocytic cells are shown in green.

**Fig 2.**
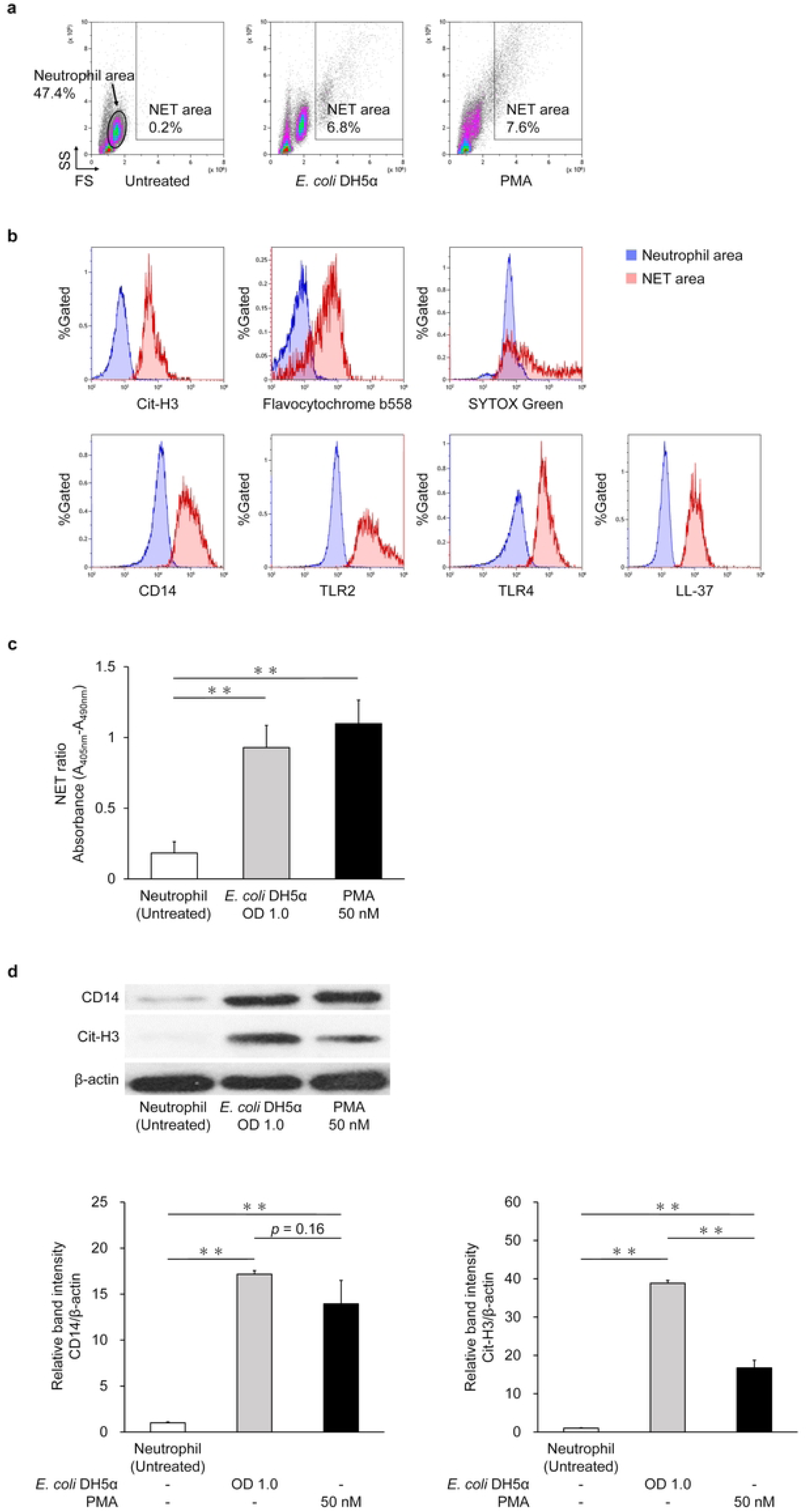
Evaluation of NET formation and associated marker expression. The percentage of the neutrophil extracellular trap (NET) area in the SS and FS plots of flow cytometry was compared among untreated neutrophils and samples following NET induction with *E. coli* DH5α or PMA. (a) Expression intensities of Cit-H3, flavocytochrome b558, SYTOX™ Green, CD14, TLR2, TLR4, and LL-37 were compared between untreated neutrophil and NET areas after *E. coli* DH5α stimulation. (b) NET ratios were compared among untreated neutrophils and samples treated with *E. coli* DH5α or PMA using enzyme-linked immunosorbent assay (ELISA). Data are presented as mean ± standard deviation (SD) (n = 3). Statistical analysis was performed using one-way analysis of variance (ANOVA) followed by Dunnett’s test. ** *p* < 0.01. (c) Western blotting was performed to examine CD14 and Cit-H3 protein levels in untreated neutrophils and those stimulated with *E. coli* DH5α or PMA. Band intensities were quantified using ImageJ and normalized to β-actin. Data are presented as mean ± standard deviation (SD) (n = 3). Data are presented as mean ± SD (n = 3). Statistical analysis was performed using one-way ANOVA followed by Tukey–Kramer’s HSD test. \*\**p* < 0.01

### NET contained high levels of CD14

The percentage of NET in areas with more than 60% Cit-H3 positivity—a hallmark of NET formation—was 6.8 ± 0.1% and 7.6 ± 0.8% following stimulation with *E. coli* DH5α and PMA, respectively, compared to 0.2 ± 0.1% in untreated cells, indicating robust NET induction by both stimulation (Fig. 2a). The MFIs of various antigens in untreated neutrophil and NET areas after *E. coli* DH5α stimulation were as follows: Cit-H3, 700 ± 13.9 and 7,679 ± 137.5; flavocytochrome b558, 619 ± 9.5 and 1,110 ± 20.4; SYTOX™ Green, 6,910 ± 28.2 and 163,045 ± 6,249.8; CD14, 12,167 ± 43.0 and 111,568 ± 1,155.4; TLR2, 9,291 ± 64.5 and 185,995 ± 151.1; TLR4, 8,572 ± 76.7 and 86,965 ± 655.8; and LL-37, 600 ± 5.3 and 11,998 ± 353.4. In NET areas stimulated by *E. coli* DH5α, fluorescence intensities of Cit-H3 and flavocytochrome b558—both critical markers of NET formation—were significantly higher than in untreated neutrophil areas, confirming NET induction (*p* < 0.01). The MFI of SYTOX™ Green, a marker of extracellular DNA, was also elevated in the NET areas, further supporting NET release. Similarly, MFI values of TLR2 and TLR4, receptors for pathogen-associated components, were increased in NET areas compared to untreated neutrophils. Furthermore, LL-37, which facilitates M1 MΦ phagocytosis, and CD14, the precursor of P-SEP, were also more abundant in the NET areas than in the untreated cells (Fig. 2b).

### Evaluation of NET Ratio by ELISA

In ELISA, the NET ratio in untreated neutrophils was 0.183 ± 0.080, whereas significantly higher ratios were observed following stimulation with *E. coli* DH5α (0.929 ± 0.157) and PMA (1.098 ± 0.166) (Fig. 2c). Western blotting revealed that the Cit-H3 protein level in untreated neutrophils was 1.000 ± 0.095, compared to 17.165 ± 0.383 and 13.940 ± 2.555 in neutrophils stimulated with *E. coli* DH5α and PMA, respectively (Fig. 2c), confirming successful NET induction. Similarly, the CD14 protein level was 1.000 ± 0.116 in untreated neutrophils, but increased to 38.845 ± 0.732 and 16.770 ± 1.967 after *E. coli* DH5α and PMA stimulation, respectively (Fig. 2d).

### M1 MΦ produce P-SEP via PR3 during NET phagocytosis

Stimulation with *E. coli* DH5α or PMA induced the release of NET into the extracellular space. Phagocytosis of NET and dead cells by M1 MΦs was also observed (Fig. 3a). Immunofluorescence staining revealed that CD68-positive M1 MΦs phagocytosed Cit-H3–positive NET and produced P-SEP intracellularly via PR3 (Fig. 3b, c). A strong correlation was observed between P-SEP levels and NET ratios measured from culture supernatants, with the regression equation y = 0.01x + 0.198 (r = 0.907) (Fig. 3d).

**Fig 3.**
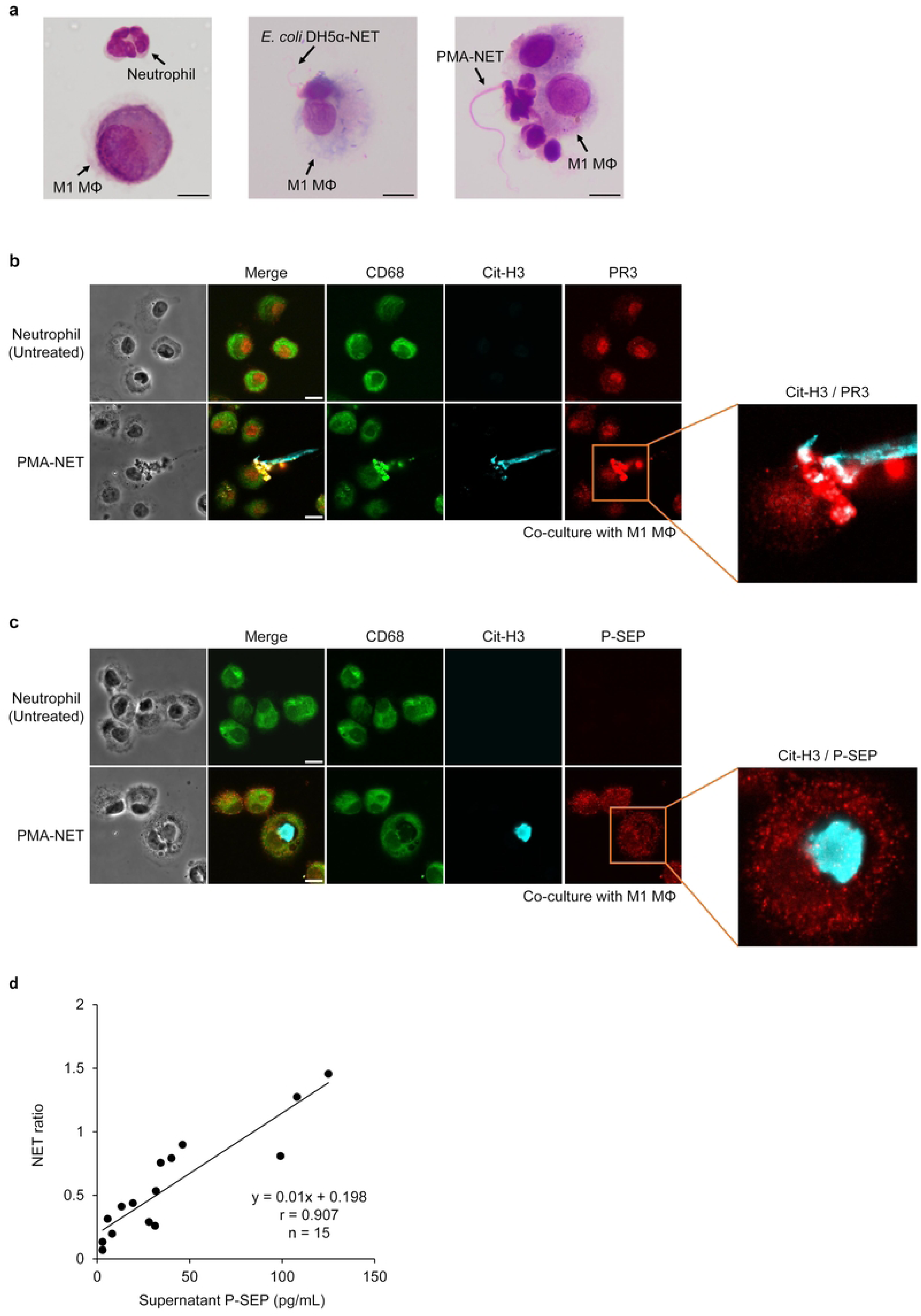
Evaluation of PR3-mediated P-SEP production during NET phagocytosis by M1 MΦ. (a) M1 MΦs were co-cultured with neutrophil extracellular trap (NET) induced by *E. coli* DH5α or PMA, followed by May–Grünwald–Giemsa (MG) staining and observation using a biological microscope. M1 MΦs were observed phagocytosing NET induced by both *E. coli* DH5α and PMA. Scale bar: 10 μm. (b, c) M1 MΦs were co-cultured with PMA-induced NET (PMA-NET), and immunofluorescence staining was performed. Confocal microscopy revealed that CD68-positive M1 MΦs phagocytosed Cit-H3–positive NET, and the intracellular localization of PR3 and presepsin (P-SEP) was examined. PR3 and P-SEP were detected in regions of M1 MΦs that had phagocytosed PMA-induced NET. Scale bar: 10 μm. (d) The concentration of P-SEP in the culture supernatant following phagocytosis of PMA-induced NET by M1 MΦs was strongly correlated with the NET ratio.

### Inhibition of PR3 reduced P-SEP production

The P-SEP level in the supernatant following co-culture of M1 MΦs and PMA-NET was 189.3 ± 17.2 pg/mL, whereas the control level, obtained from co-culture of M1 MΦs and neutrophils, was 8.8 ± 6.4 pg/mL. Inhibition of PR3 in M1 MΦs by PMSF at concentrations of 1, 10, and 50 µM resulted in concentration-dependent decreases in P-SEP production (154.7 ± 5.5, 117.7 ± 9.7, and 113.7 ± 4.0 pg/mL, respectively) (Fig. 4a). As no significant difference was observed between 10 and 50 µM (*p* = 0.99), 10 µM was selected for subsequent PR3 inhibition experiments. Similarly, treatment of M1 MΦs with elafin at concentrations of 0.1, 0.5, and 1 µM also resulted in concentration-dependent suppression of P-SEP production (100.7 ± 8.4, 84.2 ± 2.7, and 79.3 ± 3.0 pg/mL, respectively) (Fig. 4b). Since there was no significant difference between 0.5 and 1 µM elafin (*p* = 0.89), 0.5 µM was used for further PR3 inhibition assays. Overall, PR3 inhibition by 10 µM PMSF and 0.5 µM elafin in M1 MΦs co-cultured with PMA-NET reduced P-SEP levels to 117.7 ± 9.7 and 84.2 ± 2.7 pg/mL, respectively (Fig. 4c). Furthermore, western blot analysis confirmed that P-SEP protein levels in M1 MΦs that had phagocytosed NET were significantly reduced following PR3 inhibition (*p* < 0.01) (Fig. 4d).

**Fig. 4.**
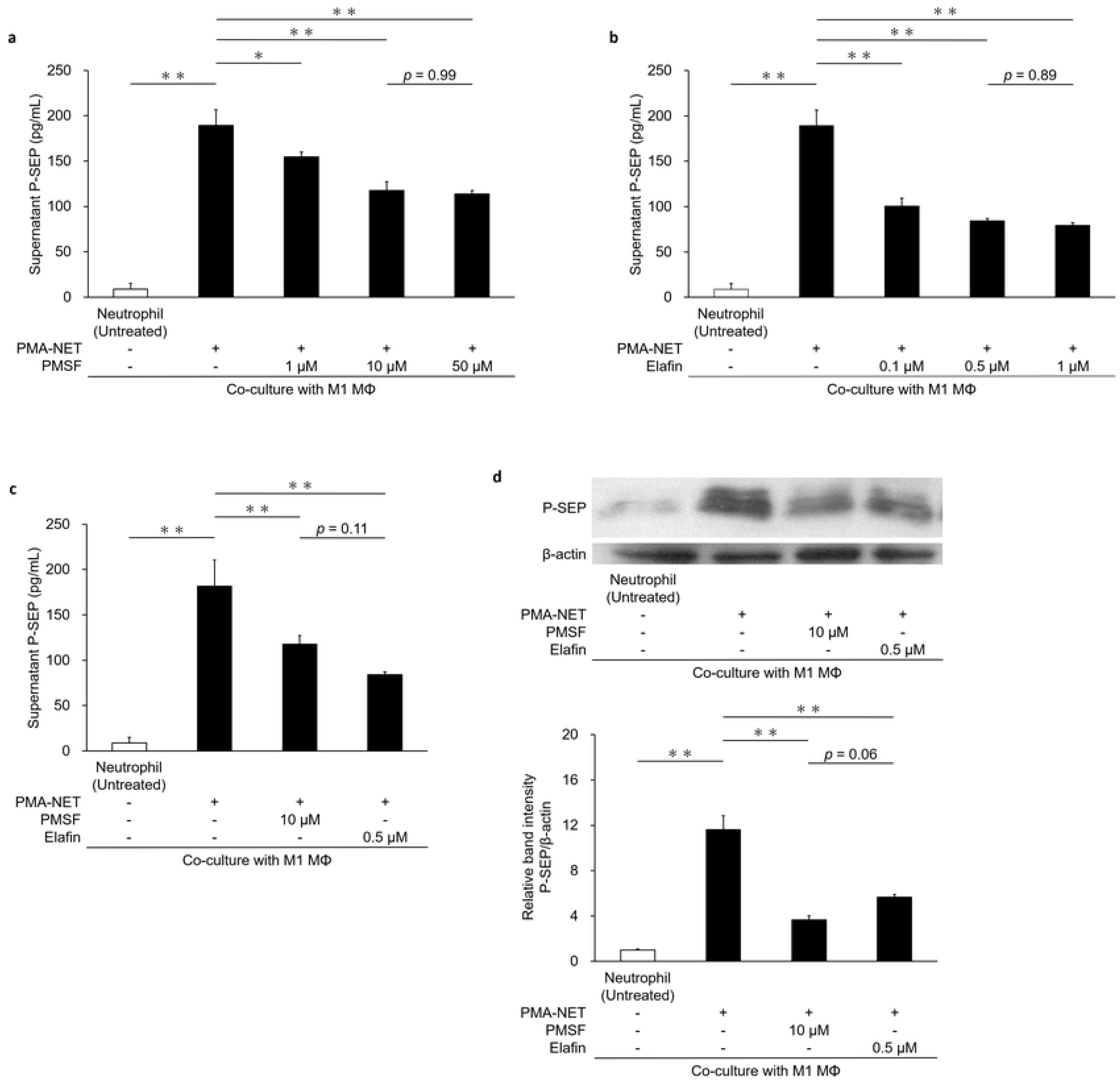
Effect of PR3 inhibition in M1 MΦs on P-SEP production during NET phagocytosis. (a, b) Supernatant presepsin (P-SEP) levels after co-culture of M1 MΦs and PMA-neutrophil extracellular trap (NET), with PMSF or elafin added at various concentrations to inhibit PR3. (c) P-SEP levels after PR3 inhibition with 10 µM PMSF or 0.5 µM elafin. (d) Western blot analysis of P-SEP protein in M1 MΦs after PR3 inhibition; protein levels were quantified using ImageJ and normalized to β-actin. Data are presented as mean ± SD (n = 3). Statistical analysis was performed using one-way analysis of variance followed by Tukey–Kramer’s HSD test. \*\**p* < 0.01, \**p* < 0.05.

### Inhibition of NET phagocytosis suppressed P-SEP production in M1 MΦ

The P-SEP level in the supernatant after co-culture of M1 MΦs with PMA-NET was significantly elevated to 200.7 ± 21.2 pg/mL, compared to 14.3 ± 2.9 pg/mL observed in co-culture with neutrophils (control) (*p* < 0.01). Subsequently, when phagocytosis was inhibited in M1 MΦs using 50 µM cytochalasin D or 10 µg/mL wortmannin, the P-SEP levels after NET co-culture decreased significantly to 49.6 ± 1.6 pg/mL and 72.3 ± 1.8 pg/mL, respectively (*p* < 0.01) (Fig. 5a). Western blot analysis further confirmed a significant reduction in P-SEP protein expression in M1 MΦs after phagocytosis inhibition, compared to M1 MΦs that had phagocytosed NET (*p* < 0.01) (Fig. 5b).

**Fig. 5.**
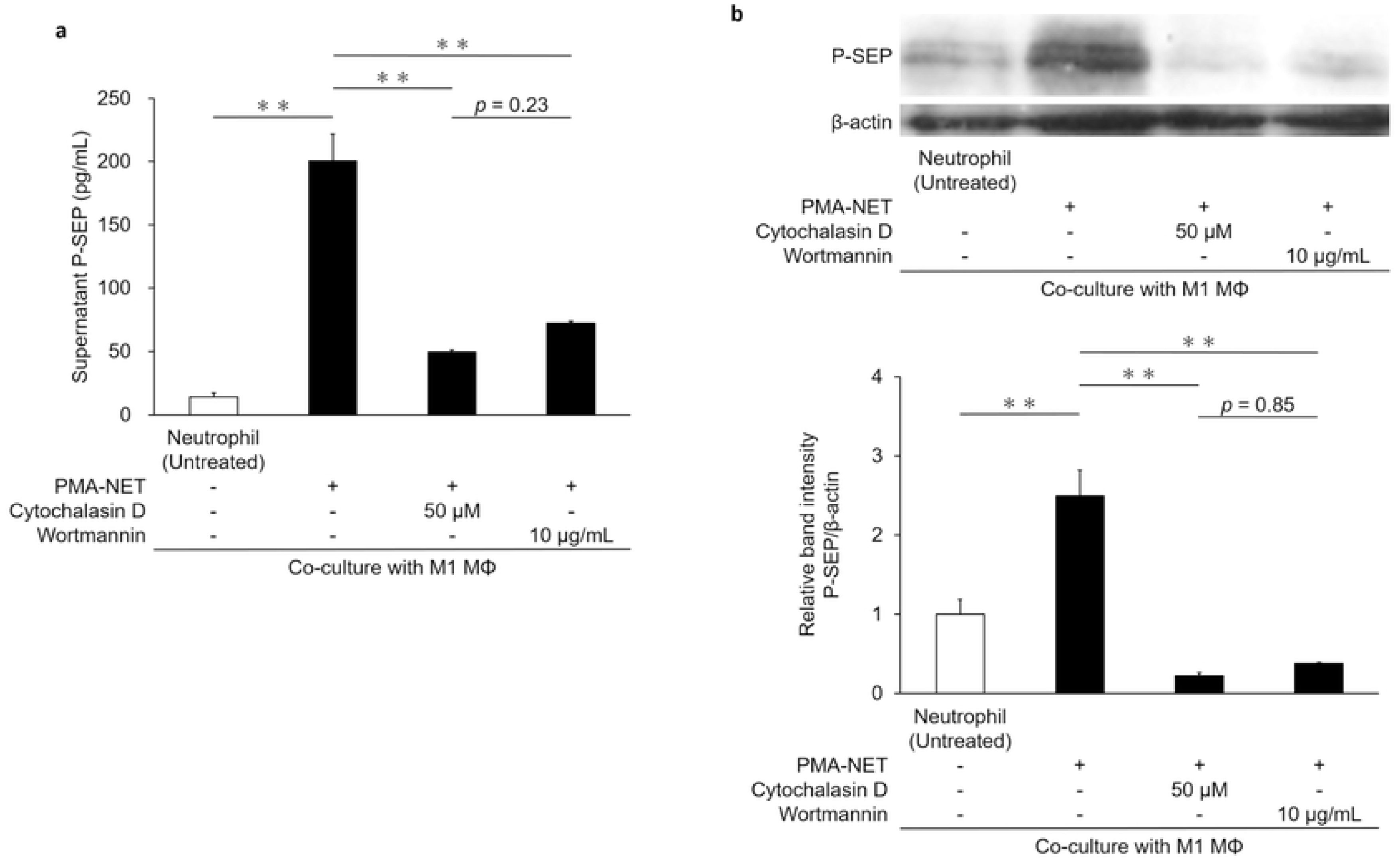
Evaluation of P-SEP production in M1 MΦs following inhibition of NET phagocytosis. (a) Supernatant presepsin (P-SEP) levels were measured after neutrophil extracellular trap (NET) co-culture with M1 MΦs treated with the phagocytosis inhibitors cytochalasin D or wortmannin. (b) Western blot analysis of P-SEP protein expression in M1 MΦs after treatment with cytochalasin D or wortmannin. Band intensities were quantified using ImageJ and normalized to β-actin. Data are presented as mean ± standard deviation (n = 3). Statistical analysis was performed using one-way analysis of variance followed by Tukey–Kramer’s HSD test. \*\**p* < 0.01, \**p* < 0.05.

### P-SEP was generated by PR3-mediated cleavage of recombinant human CD14

The P-SEP concentration in the control rCD14 solution was 7.9 ± 4.8 pg/mL, which was nearly undetectable. In contrast, the addition of PR3 markedly increased the P-SEP concentration in the supernatant to 266.3 ± 16.4 pg/mL (*p* < 0.01). In the presence of the PR3 inhibitors PMSF or elafin, the P-SEP concentration was significantly reduced to 34.8 ± 1.5 pg/mL and 41.7 ± 10.4 pg/mL, respectively (*p* < 0.01) (Fig. 6a).

**Fig. 6.**
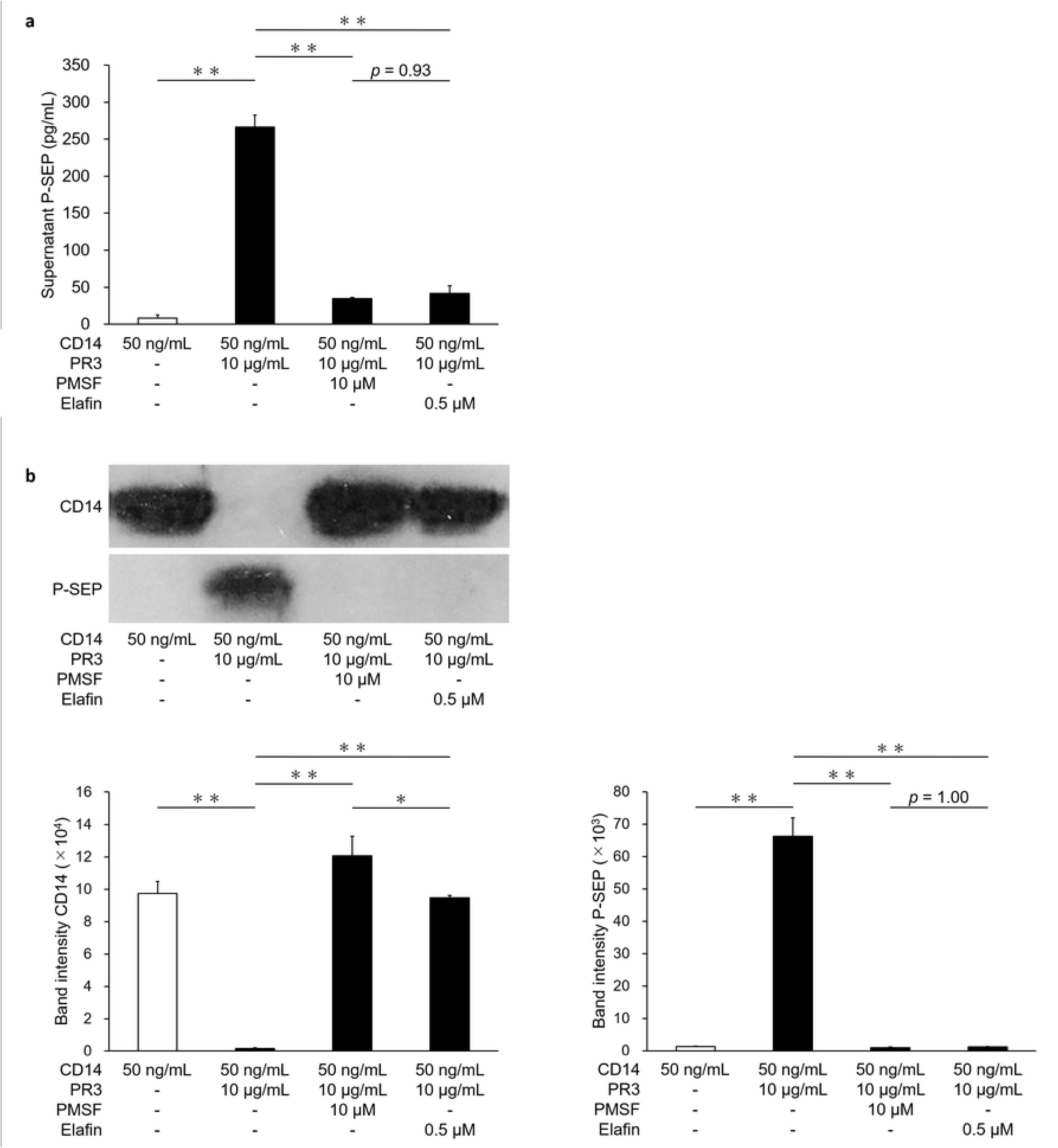
Verification of P-SEP production by addition of PR3 to recombinant human CD14. (a) Presepsin (P-SEP) levels in the supernatant were measured after addition of PR3 to rCD14. Conditions with PR3 inhibition using the inhibitors phenylmethylsulfonyl fluoride or elafin were also compared. (b) Western blot analysis of CD14 and P-SEP proteins, with band intensities quantified using ImageJ. Data are presented as mean ± SD (n = 3). Statistical analysis was performed using one-way analysis of variance followed by Tukey–Kramer’s HSD test. \*\**p* < 0.01.

### Detection of recombinant human CD14 and P-SEP proteins by western blotting

Western blot analysis of CD14 and P-SEP proteins revealed a 58 kDa band corresponding to control rCD14, with a high CD14 protein level quantified at 9.7 ± 0.7 × 10⁴ using ImageJ. However, no P-SEP band was detected, and the P-SEP protein level was low (1.3 ± 0.2 × 10³). Upon addition of PR3 to rCD14, the CD14 band disappeared due to proteolytic degradation, and the CD14 protein level decreased to 0.2 ± 0.0 × 10⁴. In contrast, a clear 13 kDa P-SEP band appeared, and the P-SEP protein level increased to 66.3 ± 5.6 × 10³. When PR3 was inhibited by PMSF or elafin, CD14 bands were retained, indicating suppression of CD14 degradation. The corresponding CD14 protein levels were 12.1 ± 1.2 × 10⁴ and 9.5 ± 0.1 × 10⁴, respectively. Under these conditions, no P-SEP bands were observed, and P-SEP protein levels remained low (0.9 ± 0.3 × 10³ and 1.2 ± 0.1 × 10³, respectively). These findings indicate that PR3 degrades CD14 to generate P-SEP, and that inhibition of PR3 suppresses both CD14 degradation and P-SEP production.

## Discussion

It has been proposed that P-SEP is generated by the degradation of CD14 by proteolytic enzymes, such as cathepsin D and elastase, in phagocytic cells. In this study, we demonstrated for the first time that PR3 also contributes to P-SEP production by degrading CD14. We first examined the expression of proteolytic enzymes in phagocytes, including M1 MΦs, Mos, and neutrophils (Fig. 1), and found that PR3 was abundantly expressed in all three cell types, similar to cathepsin D and elastase. These findings suggest that M1 MΦs—widely distributed throughout tissues—rather than circulating Mos, are the primary contributors to P-SEP production via PR3-mediated cleavage of CD14.

We then induced NET formation by stimulating neutrophils with *E. coli* DH5α or PMA and analyzed the resulting NETs using flow cytometry. To quantify NET formation, we defined the “NET area” as a region with high FS, SS, and a Cit-H3 positivity rate above 60%, a hallmark of NETs. Upon stimulation, the proportion of NETs increased to 6.8% and 7.6% for *E. coli* DH5α and PMA, respectively, compared to only 0.2% in unstimulated controls (Fig. 2a). Within the NET area, both Cit-H3 and flavocytochrome b558—an upstream component of NET formation—were upregulated. Staining with SYTOX™ Green, which labels extracellular DNA, further confirmed robust NET formation. Moreover, we observed increased expression of the bacterial receptors TLR2 and TLR4, as well as a marked upregulation of the antimicrobial peptide LL-37 (Fig. 2b). LL-37 has been reported to promote Mo-to-MΦ differentiation and to enhance macrophage phagocytic activity [29,30], suggesting that LL-37 contained in NETs may stimulate NET uptake by M1 MΦs and thereby increase P-SEP production.

More notably, flow cytometry revealed that NET areas in stimulated cultures contained higher levels of CD14—a precursor of P-SEP—than did untreated neutrophils. Given that anti-neutrophil cytoplasmic antibodies (ANCA) are known to increase CD14 expression on neutrophils and Mos [31,32], and considering the correlation observed between NET ratios and P-SEP levels in our study, it is plausible that P-SEP may also serve as a biomarker for disease activity in ANCA-associated vasculitis (Fig. 3d). Western blotting further supported this finding, as CD14 and Cit-H3 protein levels were significantly elevated in NETs induced by *E. coli* DH5α or PMA compared to untreated neutrophils (Fig. 2d), suggesting that CD14-rich NETs may be a source of P-SEP when phagocytosed by M1 MΦs (Fig. 3a).

To evaluate this mechanism, we co-cultured M1 MΦs with PMA-NETs and performed immunofluorescence staining. We confirmed that PR3 is involved in P-SEP production during phagocytosis of Cit-H3–positive NETs by M1 MΦs (Fig. 3b, c). Supernatant P-SEP levels increased significantly as M1 MΦs engulfed PMA-NETs. Inhibition of PR3 with the serine protease inhibitors PMSF and elafin led to a concentration-dependent decrease in P-SEP levels (Fig. 4a–c), and western blotting confirmed that P-SEP protein expression was reduced following treatment (Fig. 4d). Collectively, these findings indicate that M1 MΦs phagocytose NETs, internalize NET-derived CD14, and degrade it via intracellular PR3 to generate P-SEP.

To further validate that P-SEP production is driven by NET phagocytosis, we inhibited M1 MΦ phagocytic function using cytochalasin D and wortmannin. As shown in our previous study, cytochalasin D suppressed M1 MΦ phagocytosis and reduced P-SEP production in a concentration-dependent manner. In the present study, we demonstrated that wortmannin also effectively inhibited phagocytosis, leading to a similar dose-dependent reduction in P-SEP levels (Supplementary Fig. 3). Western blotting again confirmed that inhibition of phagocytosis led to decreased P-SEP production (Fig. 5a, b), supporting the notion that M1 MΦs produce P-SEP by internalizing and processing CD14-rich NETs via PR3. Although these data support the role of PR3 in CD14 degradation, it should be noted that selective PR3 inhibitors are currently unavailable. In this study, we utilized PMSF, which broadly inhibits serine proteases including PR3, and elafin, which inhibits both PR3 and elastase [35]. As these agents are not PR3-specific, their use alone cannot conclusively prove that PR3 is solely responsible for CD14 cleavage and P-SEP generation.

Yard et al. previously reported that PR3 degrades rCD14 [34]. In our study, we confirmed that addition of PR3 to rCD14 in vitro led to CD14 degradation and production of the 13 kDa presepsin. While the P-SEP level in the control rCD14 solution was low (7.9 ± 4.8 pg/mL), the addition of PR3 increased it dramatically to 266.3 ± 16.4 pg/mL (Fig. 6a, b), clearly demonstrating that PR3 can directly generate P-SEP through CD14 cleavage.

We propose that *in vivo*, P-SEP may be produced by PR3 released into plasma in association with NETs, which then degrades membrane-bound CD14 on phagocytes and soluble CD14 in circulation. In addition to their antimicrobial phagocytic function, neutrophils contribute to host defense by releasing NETs into the extracellular space to trap and neutralize pathogens. Importantly, NETs were shown to contain significantly higher amounts of CD14 than resting neutrophils. In this study, we demonstrated that tissue-resident M1 MΦs phagocytose NETs, and that PR3 within these cells degrades internalized CD14 to produce P-SEP (Fig. 7). As P-SEP levels positively correlated with NET abundance, its measurement may reflect NET activity *in vivo*.

**Fig. 7.**
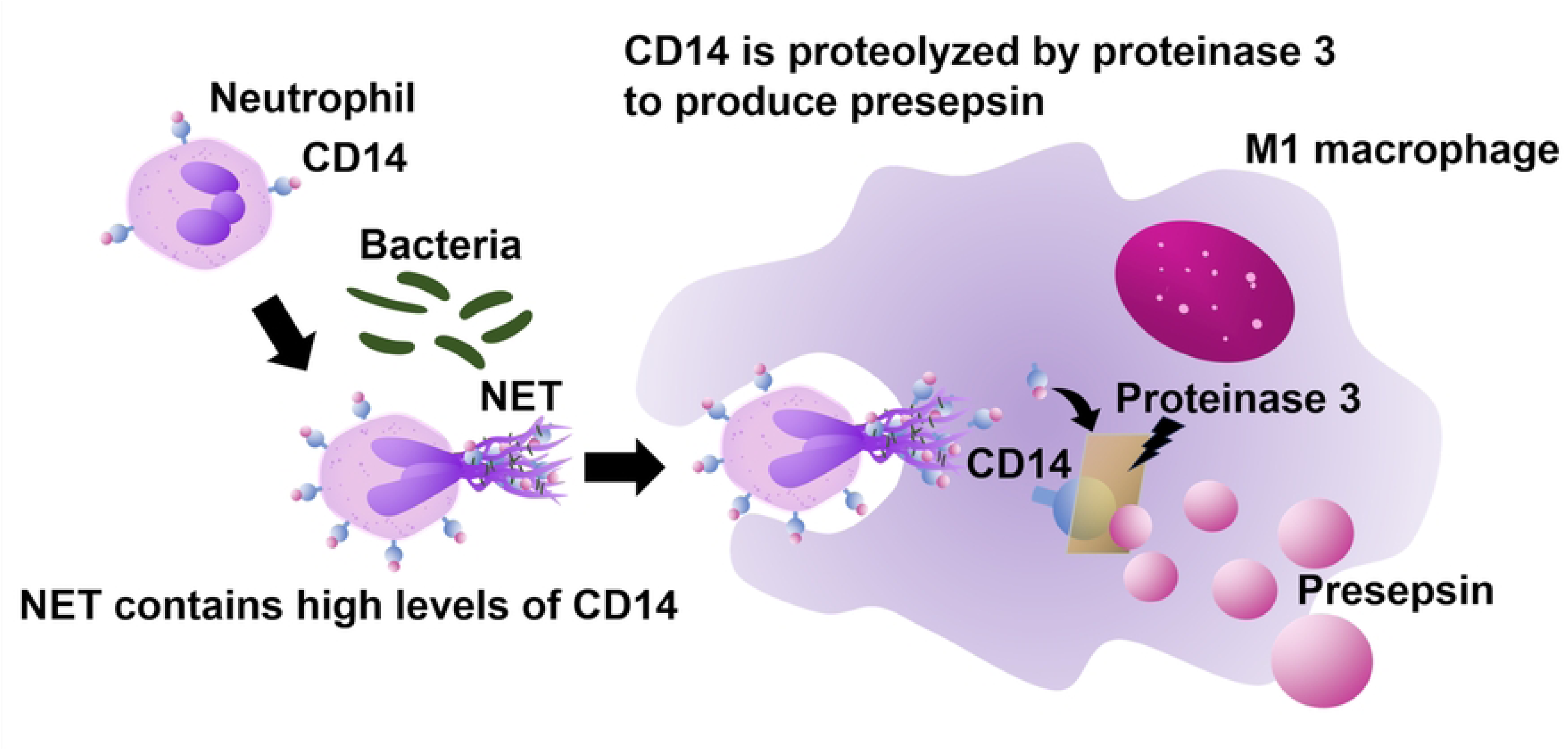
Schematic illustration of the mechanism by which M1 MΦs produce P-SEP via PR3 following NET phagocytosis. Upon bacterial infection of blood vessels, neutrophils release neutrophil extracellular traps (NETs) as a host defense mechanism. M1 MΦs phagocytose CD14-rich NETs, and the engulfed CD14 is degraded by the proteolytic enzyme PR3 within the macrophages, resulting in the generation of presepsin (P-SEP). Subsequently, P-SEP is released extracellularly from M1 MΦs, leading to elevated blood levels of P-SEP.

Thus, beyond its established role as a sepsis biomarker, P-SEP holds promise as a novel marker of disease activity in autoimmune diseases characterized by increased NET formation. These findings provide a foundation for future clinical investigations aimed at validating P-SEP as a diagnostic and monitoring tool for NET-related pathologies.

## Acknowledgments

The authors thank Kamon Shirakawa of PHC Holdings Corporation (Tokyo, Japan) for productive discussions during this project. In addition, the authors thank all blood donors who allowed us to conduct experiments during this study. We also thank Editage (www.editage.com) for English language editing.

## Data Availability Statement

Data used and analyzed during the current study are available from the corresponding author upon reasonable request.

## Supplementary material

Supplemental material to this article is available online.

## Figure Captions

**Supplementary Fig. 1. Unprocessed images of the western blot gels shown in Fig. 2d**

Western blot analysis of CD14, Cit-H3, and β-actin expression in NETs induced by 50 nM PMA or OD 1.0 *E. coli* DH5α. The regions enclosed by white lines were used in the preparation of Fig. 2d.

**Supplementary Fig. 2. Unprocessed images of the western blot gels shown in** Fig. 4d

Western blotting was performed to detect β-actin and P-SEP in samples co-cultured with M1 MΦs and neutrophils, with PMA-NETs, and with each PR3 inhibitor. The area enclosed by the white line was used in Fig. 4d.

**Supplementary Fig. 3. Effect of wortmannin on P-SEP production by inhibiting NET phagocytosis in M1 MΦs**

Supernatant P-SEP levels were measured after co-culture of M1 MΦs with PMA-NETs in the presence of wortmannin. Inhibition of NET phagocytosis suppressed P-SEP production. Data are presented as mean ± SD (n = 3). Statistical analysis was performed using one-way ANOVA followed by Tukey–Kramer’s HSD test. ** *p* < 0.01.

**Supplementary Fig. 4. Unprocessed images of the western blot gels shown in** Fig. 5b

Western blot analysis of β-actin and P-SEP expression in M1 MΦs co-cultured with neutrophils, PMA-NETs, or phagocytosis inhibitors. The area enclosed by the white line was used in Fig. 5b.

**Supplementary Fig. 5. Unprocessed images of the western blot gels shown in** Fig. 6b

Western blot detection of CD14 and P-SEP in samples treated with PR3 or PR3 plus inhibitors in recombinant CD14 (rCD14) solution. The area enclosed by the white line was used in Fig. 6b.

## Notes

### Competing Interest Statement

The authors have declared no competing interest.

